# Contrasting patterns of subtelomeric satellite superfamily in the *Cannabaceae* family

**DOI:** 10.1101/2025.01.31.635892

**Authors:** Lucie Horáková, Václav Bačovský, Bohuslav Janoušek, Josef Patzak, Roman Hobza

**Affiliations:** Department of Plant Developmental Genetics, Institute of Biophysics of the Czech Academy of Sciences, Kralovopolska 135, 612 00 Brno, Czech Republic; Department of Experimental Biology, Faculty of Science, Masaryk University, Kamenice 5, 625 00 Brno, Czech Republic; Hop Research Institute Co. Ltd, Kadaňská 2525, 438 46 Žatec, Czech Republic

**Keywords:** *Humulus*, subtelomeric DNA, sex chromosomes, satellite DNA

## Abstract

Satellite DNA (satDNA) is a rapidly evolving component of plant genomes, often found in centromeric, telomeric, and heterochromatic regions. Due to their variability and species- or population-specific distribution, satDNA serves as valuable cytogenetic markers for studying chromosomal rearrangements and karyotype evolution in closely related species. In dioecious species *Cannabis sativa, Humulus lupulus*, and *Humulus japonicus*, previous studies have identified species-specific subtelomeric repeats CS-1, HSR1, and HJSR. While these satellites have been used to differentiate sex chromosomes from autosomes, their evolutionary origins, sequence variation and pattern of conservation among related species remain largely unexplored. In this study, we combine bioinformatics analysis with molecular cloning to analyze sequence similarity among these shared satellites and determine their inter-specific chromosomal localization using fluorescence *in situ* hybridization (FISH). Our results reveal that HSR1 and HJSR satellites are shared among all studied species suggested oaring from a common ancestor. In contrast, CS-1 satellite exhibit higher sequence divergence. Although all satellites are predominantly localized in subtelomeric regions, CS-1 in *H. lupulus* and HSR in *C. sativa* are localized in pericentromeric regions. These findings provide new insight into the evolutionary dynamics of satDNA in *Cannabaceae* family and its role in genome organization.

## 1. Introduction

Plant genomes are predominantly composed of repetitive DNA, which can constitute up to 85% of the total genome (Novák et al., 2020). The repetitive fraction of a genome is classified into two major categories: dispersed repeats (transposable elements) and tandem repeats (including satellite DNA, satDNA). SatDNA is composed of monomer sequences arranged in long tandem arrays, accumulated primarily in chromosomal subdomains characterized by a compact and heterochromatic organization, such as (peri)centromeres and (sub)telomeres (Garrido-Ramos, 2015). SatDNA represents one of the most rapidly evolving components of eukaryotic genomes, differing among related species in (i) nucleotide sequence, (ii) copy number, (iii) length of monomer subunits, and (iv) chromosome localization (Plohl et al., 2008; Lower et al., 2018). SatDNA is often enriched within specialized genomic elements (e.g. sex chromosomes), which may accumulate large numbers of satellite families due to low or suppressed recombination (Palacios-Gimenez et al., 2017; Jesionek et al., 2021). In addition to other factors, these large satDNA arrays have a role in chromosome evolution and segregation, highlighting their functional significance (Plohl et al., 2012). The repetitive nature of satDNA makes it a robust and powerful tool for cytogenetic studies, enabling detailed analysis of individual chromosomes (Hobza et al., 2024), chromosomal rearrangements, and evolutionary dynamics among related species (Jagannathan et al., 2017; Schmidt et al., 2019, 2024). According to the library hypothesis, related species share a set of satellite sequences with a certain level of sequence divergence and variation in abundance. Differential amplification of specific satellite sequences leads to unique satDNA collections in closely related species, which display species- or population-specific profiles (Fry et al., 1973; Mestrovic et al., 1998; Plohl et al., 2012). This hypothesis has been extensively studied in various plants (Koukalova et al., 2010; Belyayev et al., 2019, 2020), fish (Utsunomia et al., 2017; Goes et al., 2022), and other eukaryotic organisms (Mestrovic et al., 1998; Camacho et al., 2022). Furthermore, satDNA monomers are typically homogenized within a species as a consequence of concerted evolution, a process that maintains sequence uniformity within a species while promoting interspecies divergence (Waye and Willard, 1989; Durfy and Willard, 1990; Wei et al., 2014).

The *Cannabaceae* family includes the genera *Cannabis* and *Humulus*, along with eight additional genera widely distributed across tropical and temperate regions (Yang et al., 2013; Jin et al., 2020). The genus *Cannabis*, thought to have originated in East Asia, consist of a diploid species, *Cannabis sativa* (Lapierre et al., 2023), which is used for fiber, oil, or as euphoric intoxicant and therapeutic drug (Small, 2015). The sister genus *Humulus* includes three species: *Humulus lupulus* L., *H. japonicus* Siebold & Zucc. (synonym *H. scandens* (Lour.) Merr.), and Chinese endemic species *H. yunnanensis* Hu. (Ling and Zhang, 2019). *H. lupulus* is essential for beer brewing and together with *H. japonicus*, contains unique secondary metabolites with strong therapeutic potential (Yu et al., 2007; Zanoli and Zavatti, 2008; Ryu et al., 2017; Jiang et al., 2018). Divergence between *C. sativa* and *H. lupulus* occurred between 16 – 27.8 mya (McPartland, 2018; Jin et al., 2020; Prentout et al., 2021; Padgitt-Cobb et al., 2023), whereas *H. lupulus* and *H. japonicus* diverged relatively recently, around 3.7 – 10.7 mya (Murakami, 2000; Murakami et al., 2006; Jin et al., 2020). Members of the *Cannabis* and *Humulus* genera are primarily dioecious, with monoecious cultivars occurring rarely. It is hypothesized that dioecy likely evolved before the divergence of these two genera, at least 21-25 mya (Prentout et al., 2021).

The three species within the *Cannabis* and *Humulus* genera, namely *C. sativa, H. lupulus* and *H. japonicus*, exhibit significant differences in genome size, ranging from 808 Mb in *C. sativa* (Gao et al., 2020) to 1.8 Gb in *H. japonicus* (Grabowska-Joachimiak et al., 2006) and 2.8 Gb in *H. lupulus* (Padgitt-Cobb et al., 2023). A substantial proportion of these genomes consist of repetitive DNA: 64-74.8% in *C. sativa* (Pisupati et al., 2018; Gao et al., 2020; Lynch et al., 2024), 61.3-64.5% in *H. lupulus* (Pisupati et al., 2018; Padgitt-Cobb et al., 2023; Horáková et al., 2024), and 66.8% in *H. japonicus* (Zhang et al., 2023). In line with patterns observed in plant genomes (Feschotte et al., 2002), the long terminal repeat retrotransposons (LTR-RTs) are the most abundant class of repetitive DNA across *Cannabis* and *Humulus* species (Pisupati et al., 2018; Zhang et al., 2023; Lynch et al., 2024). A burst of LTR-RT proliferation may have contributed to the X chromosomes enlargement and rearrangements in both *Humulus* species (Akagi et al., 2024). Interestingly, the *H. japonicus* sex chromosomes underwent X-autosome fusion that involved autosome 3, creating neo-Y chromosomes, allocating at least one terminal end of A3 to end of the X chromosome (Akagi et al., 2024). In contrast, the accumulation of Ty1/*Copia* solo-LTRs has predominantly shaped the Y chromosome in *C. sativa* (Lynch et al., 2024). Meanwhile, satDNA constitutes only a small fraction of the repetitive DNA in *Humulus* species, accounting for 0.3-2% of the genome (Zhang et al., 2023; Horáková et al., 2024). Recent studies using long-read PacBio sequencing and advanced bioinformatics tools have characterized tandem repeat families in *H. lupulus* (Easterling et al., 2020). This approach has led to the identification of the most abundant DNA repeats in *H. japonicus* (Zhang et al., 2023). To date, the most valuable cytogenetic markers for identifying autosomes and sex chromosome in *Cannabis* and *Humulus* species include three members of putative superfamily cluster CS-1 (*Cannabis sativa* 1; Divashuk et al. 2014), HSR1 (*Humulus* subtelomeric repeat 1; Divashuk et al. 2011), and HJSR (*Humulus japonicus* subtelomeric repeat; Alexandrov et al. 2012). Both subtelomeric HSR1 and HJSR satellites were identified previously using DNA digestion with the *Kpn*I restriction endonuclease (Divashuk et al., 2011; Alexandrov et al., 2012), while CS-1 was derived from HSR1 as a repeat with low homology to the HSR1 satellite (Divashuk et al., 2014). This may suggest common origin for both repeats, classifying them into one superfamily clusters as they are distributed across different regions in the genomes as observed in other plats species as described in *Rumex acetosa* (Navajas-Pérez et al., 2006, 2009b; Cuñado et al., 2007; Quesada Del Bosque et al., 2011). Despite the independent identification of these subtelomeric satellites, they exhibit further remarkable similarity across both *Humulus* and *C. sativa* species, displaying either subtelomeric or pericentromeric position or sequence structure. Nevertheless, the origin and the evolution of these large satellite arrays remain poorly understood, particularly compared to those described in other plants.

In this study, we isolated and compared superfamily clusters represented by three tandemly arrayed repeats HSR1 (*H. lupulus*-specific), HJSR (*H. japonicus*-specific), and CS-1 (*C. sativa*-specific). We identified their interspecific localization using fluorescent *in situ* hybridization (FISH) on metaphase chromosomes in both sexes and quantified their genome abundance, sequence similarity, and genomic variability combining short Illumina reads and bioinformatic approaches together with molecular cloning, within each species. Given the parallel origin of HSR1, HJSR, and CS-1, we aimed to compare (i) the genetic differentiation of the HSR1 and HJSR satellites in the context of *H. japonicus* speciation, and (ii) the origin and divergence of this subtelomeric satellite family. We also discussed the sequence similarity and chromosomal positioning of the HSR1 and CS-1 satellites.

## 2. Materials and methods

### 2.1. Plant material

Female *Humulu*s *lupulus* cv Saaz hop (Osvald’s clone 72) and male *H. lupulus* Lib male (15181) (2n = 18 + XX/XY) were provided by the Hop Research Institute Co. Ltd. in Žatec (Czech Republic). Female and male *Humulus japonicus* (2n = 14 + XX/XY_1_Y_2_) and *Cannabis sativa* cv Kompolti (2n = 18 + XX/XY) plants were grown from seeds obtained from W. Legutko (Poland) and SEMO a.s. (Czech Republic), respectively. All plants were grown in a greenhouse under controlled conditions (16h daylight/ 8h dark photoperiod) at the Department of Plant Developmental Genetics in Brno (Czech Republic). The sex of *H. japonicus* plants was determined based on floral morphology and chromosome number. In *C. sativa* and *H. lupulus*, sex was determined using PCR with male-specific molecular markers: MADC2 for *C. sativa* (Mandolino et al., 1999) and OPJ9 for *H. lupulus* (Polley et al., 1997). Both male-specific markers were amplified using Taq polymerase (Top Bio) according to the manufacturer’s instructions. PCR cycling conditions followed protocols described by Razumova et al. (2016) and Patzak et al. (2002). Male and female plants of each species were selected for further analysis based on the PCR results (Supplementary Figure S1).

### 2.2. Isolation of DNA and Low coverage genome sequencing

Genomic DNA was extracted from young leaves of male *C. sativa, H. lupulus*, and *H. japonicus*, using the NucleoSpin Plant II (Macherey-Nagel GmbH and Co. KG., Germany), following the manufacturer’s protocol. Libraries were prepared using the NEBNext® Ultra™ II DNA Library Prep Kit. Genomic DNA from male and female plants of *C. sativa* and *H. japonicus* was low coverage sequenced using an Illumina MiSeq sequencer, generating 300 bp paired-end-reads at the Centre of Plant Structural and Functional Genomics (Olomouc, Czech Republic). Additionally, we used the library of *H. lupulus* described in Horáková et al. (2024).

### 2.3. Analysis of repeats using Repeat Explorer

The FastQC tool (available at http://www.bioinformatics.babraham.ac.uk/projects/fastqc) was used to assess the quality of sequencing reads. Furthermore, reads were pre-processed based on quality (Q30) with subsequent adaptor trimming, filtering out short or unpaired sequences, and all reads were trimmed to a uniform length of 200 bp using Trimmomatic 0.32 (Bolger et al., 2014). To identify repetitive DNA composition, these datasets were analyzed using RepeatExplorer2 (Novák et al., 2010, 2013). The clustering of randomly selected 2 x 500 000 reads was performed by default setting.

### 2.4. Ligation of PCR products and cloning

The major satellites including CS-1 (GenBank accession number JX402748.2), HSR1 (*Humulus* subtelomeric repeat, GenBank accession number GU831574.1), HJSR (*Humulus japonicus* subtelomeric repeat, GenBank accession number GU831573.1), and 45S rDNA were amplified by PCR using specific primers (Supplementary Table S1) and Q5 High-Fidelity DNA polymerase (M0491S; NEB), following manufacturer’s instructions. PCR cycling conditions were as follows: 95°C for 4 min followed by 35 cycles of 94°C for 30 s, 55°C for 35 s, 72°C for 30 s, and a final extension step at 72°C for 10 min. The selected units for each satellite (Supplementary Figure S2) were extracted and purified from the gel using the QIAquick Gel Extraction Kit (QIAGEN GmbH -Hilden, Germany). Purified fragments (Supplementary Figure S2) were individually cloned into the pJET1.2 vector using the CloneJET PCR Cloning Kit (K1231; ThermoFisher) following the manufacturer’s instructions. The ligation reactions were incubated overnight at 16 °C and subsequently desalted. Competent *E. coli* cells were transformed with ligation reactions via electroporation using the MicroPulser Electroporator (Bio-Rad). After 30 min of incubation, the transformed cells were spread on Petri dishes with solid LB medium containing ampicillin (100 mg/L) and incubated overnight at 37 °C. The next day, colony PCR was performed to screen for positive clones. Individual bacterial colonies were used as templates for amplification with pJET1.2-specific primers. Primers and nucleotides were removed using ExoSAP reaction and sequenced in Macrogen (Amsterdam, Netherlands).

### 2.5. DNA probes preparation

Purified PCR products (1µg) from each satellite and species obtained from the gel as described above were labeled with Atto488 NT (PP-305L-488) and Atto550 NT (PP-305L-550) using Nick Translation Labelling kits (Jena Bioscience, Germany). The labeling reactions were incubated at 15°C for 90 min. Labeled DNA probes were directly used in hybridization mixture for FISH. A telomeric repeat (Arabidopsis-type, TTTAGGG) was amplified according to Ljdo et al. (1991), without template DNA. PCR cycling conditions were as follows: 95°C for 4 min followed by 35 cycles of 94°C for 30 s, 55-60°C for 35 s, 72°C for 30 s and final extension step at 72°C for 10 min.

### 2.6. Mitotic chromosome preparation and Fluorescence *in situ* hybridization

Mitotic chromosomes were prepared from young leaves (2-5 mm in length) of *C. sativa, H. lupulus*, and *H. japonicus* as described in Horáková et al. (2024). FISH was performed on mitotic metaphase chromosomes according to Sacchi et al. (2024) under two stringency conditions (77% and 68%; Supplementary Table S2). FISH with 77% stringency was used to confirm the subtelomeric localization of CS-1 in *C. sativa*, HSR1 in *H. lupulus*, HJSR in *H. japonicus*, and 45S rDNA. Low-stringency FISH (68%) was used to determine the interspecific localization of satellites. Chromosomes were counterstained with 4′,6′-diamidino-2-phenylindole (DAPI) in Vectashield Antifade Mounting Medium. Images were captured using an Olympus AX70 epifluorescence microscope equipped with a CCD camera and processed using Adobe Photoshop. FISH experiments were performed in triplicate, with at least ten metaphases analyzed per experiment.

### 2.7. Phylogenetic analysis of subtelomeric satellites

The sequences were aligned in MAFFT in two rounds. At first, the adjusted direction option was used to ensure the correct orientation of sequences. The alignment was further improved in MAFFT using an iterative refinement method incorporating local pairwise alignment (mafft-linsi). The phylogenetic tree was constructed using maximum-likelihood approach using IQ-TREE. The substitution model (HKY+F+G4) was chosen in IQ-TREE based on the BIC criterion. The support values were obtained using the ultrafast bootstrap method (with 1000 replicates).

To obtain sequence logos for each satellite-species combination, we mapped the reads from a given species to the reference, which was represented by a 60% consensus sequence prepared from sequenced clones of PCR products obtained in the species in which that satellite was originally described (for primers, see Supplementary Table S1). In several satellite-species combinations, low-coverage sequencing did not allow sufficient coverage of the reference, so the sequence logo was prepared based on the cloned PCR products. DNA reads were trimmed using Trimmomatic (Bolger et al., 2014) and mapped to the reference using BWA-MEM (Li, 2013). SAM files were converted to fasta alignments using a simple Python script (sam2fasta.py; https://sourceforge.net/projects/sam2fasta/). Sequence logos were generated from fasta alignments using a locally installed version of the Weblogo 3 program (Crooks et al., 2004) with sequence-type set to ‘dna’.

## 3. Results

### 3.1. Comparison of major satellite distribution among the related species

To analyze the interspecific distribution of previously identified satellites – HSR1 (Divashuk et al., 2011), HJSR (Alexandrov et al., 2012), and CS-1 (Divashuk et al., 2014), we performed FISH on metaphase chromosomes of closely related species: *C. sativa, H. lupulus*, and *H. japonicus* (Figure1 and Table 1).

**Figure 1.**
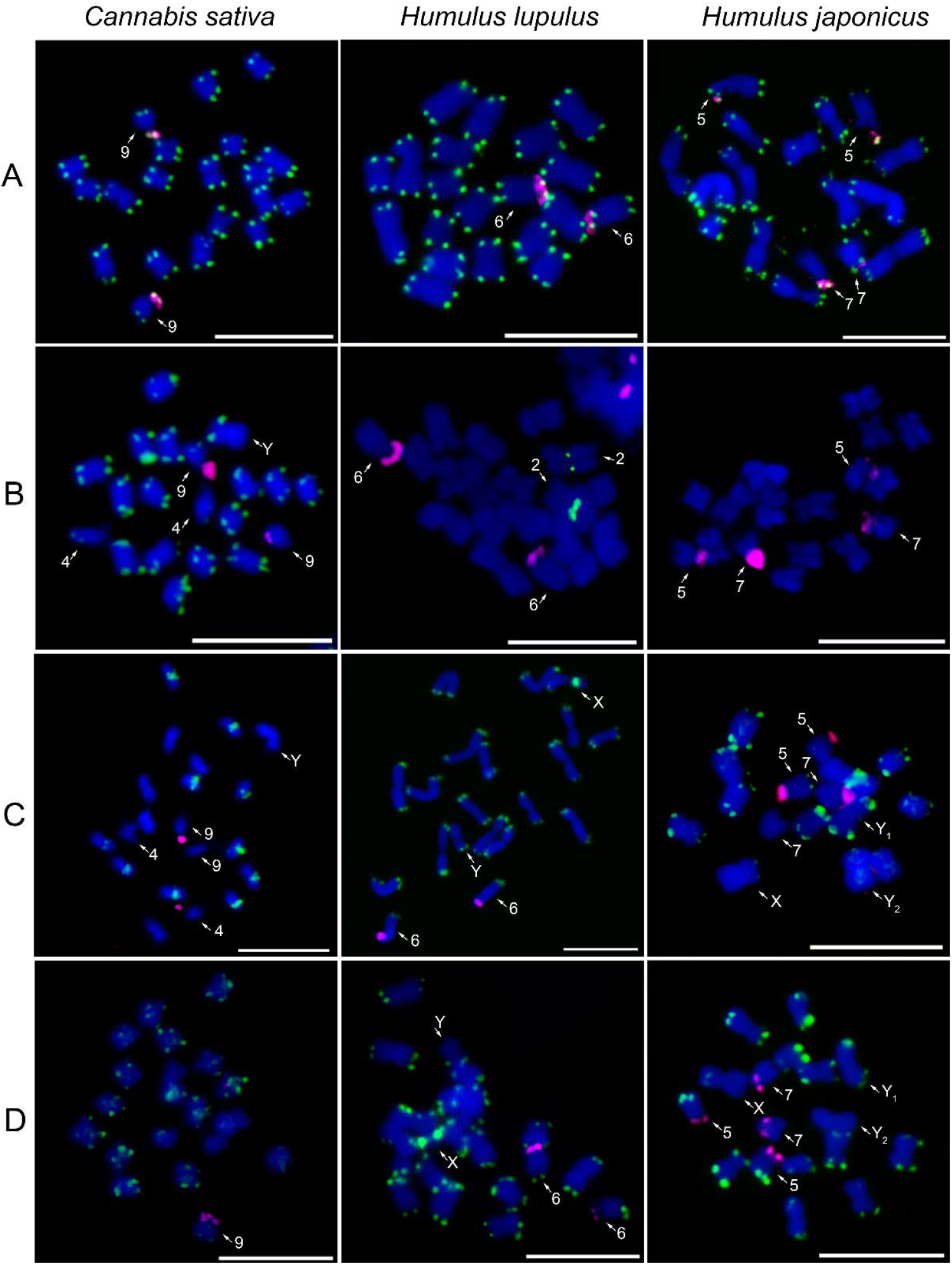
Chromosomal distribution of major satellites in studied species. The localization of (A) the telomeric sequence motif TTTAGGG (green), (B) the CS-1 satellite (green), (C) the HSR1 satellite (green), (D) the HJSR satellite (green), and 45S rDNA (magenta) on male mitotic metaphase chromosomes of *Cannabis sativa, Humulus lupulus*, and *H. japonicus*. Note the reduced number of CS-1 positive regions in *H. lupulus* and the absence of this satellite in *H. japonicus* (B). Interestingly, HSR1 is localized within the (peri)centromeres in *C. sativa*, while it displays a subtelomeric position in both *H. lupulus* and *H. japonicus* (C). Mitotic chromosomes were counterstained with DAPI. Arrows indicate differentiated autosomes and sex chromosomes. Scale bar = 10 µm.

**Table 1.**
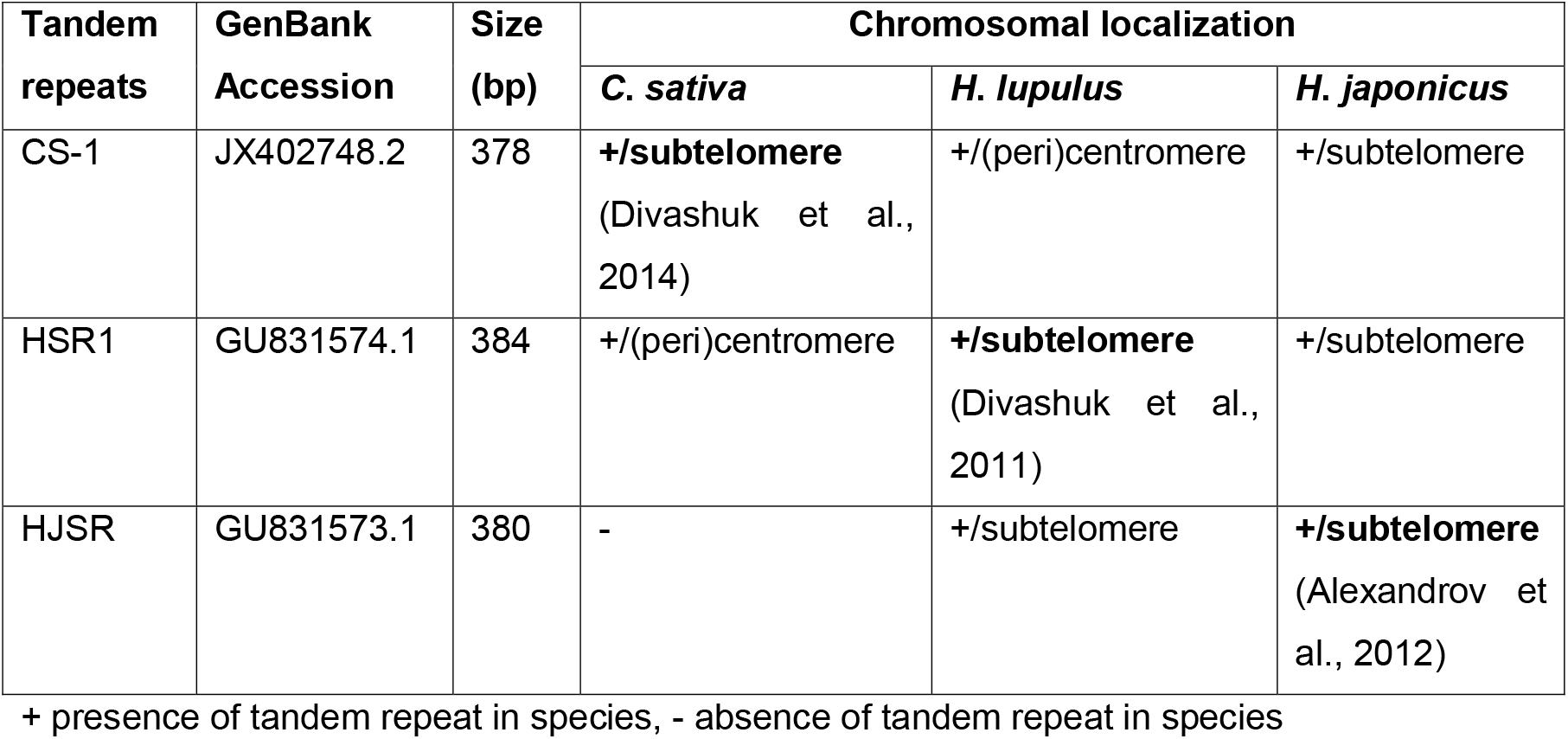
Analyzed tandem repeat in *C. sativa, H. lupulus, H. japonicus*, their size and distribution patterns.

Additionally, *Arabidopsi*s-type telomeric (TTTAGGG) and 45S rDNA probes were used to label chromosome ends and enable the identification of previously described chromosomes (Figure 1A). Telomeric signals are present at the terminal regions of all chromosomes, without any interstitial telomeric signals (Figure 1A and Supplementary Figure S3A). The distribution of 45S rDNA varies among studied species, with localization on chromosome 9 in *C. sativa*, chromosome 6 in *H. lupulus*, and two pairs of autosomes in *H. japonicus*, specifically chromosomes 5 and 7 (Figure 1A).

We observed chromosome-specific differences in the localization of major satellites. In *C. sativa*, CS-1 is localized in the subtelomeric regions of both arms of all autosomes, except for the q-arm of chromosome 4, and p-arm of chromosome 9, which is positive for 45S rDNA. The Y chromosome displays the same absence of CS-1 on the q-arm within the karyotype (Figure 1B). This distinct distribution of CS-1, along with the Y chromosome size and DAPI banding pattern, facilitates its precise identification (Figures 1B-C). However, the identification of X chromosome is still difficult, based only on CS-1 distribution and factors mentioned above (Figures 1B-C). In *H. lupulus*, the CS-1 satellite is located in the (peri)centromeric region of chromosome 2 (Figure 1B) simultaneously with 5S rDNA in the subtelomeric region (Supplementary Figure S4). Although the CS-1 satellite was amplified with the designed primers in *H. japonicus* (Supplementary Figure S2), we did not observe any clear signals on metaphase chromosomes of either sex (Figure 1B and Supplementary Figure S3B).

The *H. lupulus*-specific repeat, HSR1 satellite, is localized in the (peri)centromeric regions of ten chromosomes in males (Figure 1C) and on eleven chromosomes in females *C. sativa* (Supplementary Figure S3C). The observed structural chromosomal heterozygosity suggests the hybrid origin of female *C. sativa* cv Kompolti or a possible localization of HSR1 on one of the X chromosomes. The simultaneous hybridization of 45S rDNA, HSR1, and CS-1 satellites shows that the HSR1 satellite is absent from chromosomes 4, 9, and Y (Supplementary Figure S5). In contrast, the HSR1 satellite is localized in the subtelomeric regions of almost all autosomes of *H. lupulus*, except for the p-arm of chromosome 6, which carries 45S rDNA, the p-arm of X chromosome, and the q-arm of chromosome Y (Figure 1C). Notably, the X chromosome exhibits a strong HSR1 signal in the pericentromeric region of its p-arm. In *H. japonicus*, the HSR1 probe is distributed in the subtelomeric regions of almost all chromosomes (Figure 1C), showing similarities to the HJSR probe (Figure 1D). However, one chromosome pair and the chromosome Y_2_ are absent or strongly underrepresented form HSR1.

In *C. sativa*, HJSR is found mainly in subtelomeric regions, although, some chromosomes lack the HJSR signal (Figure 1D). The distribution of the HJSR satellite in *H. lupulus* was identical to that of the HSR1 satellite, including its localization on the sex chromosomes (Figure 1D and Supplementary Figure S3D). In *H. japonicus*, the HJSR satellite is present in subtelomeric regions, except for one arm of X chromosome, p-arm of chromosome 5, and both arms of chromosome 7, which carry 45S rDNA. No HJSR signal was observed on the Y_2_ chromosome (Figure 1D). The distribution of 45S rDNA across all analyzed species aligns with previous studies. The distribution of CS-1, HSR1 and HJSR satellites in female plants are shown in Supplementary Figure S3. We observed no sex-specific differences in the distribution of these satellites (Figure 1 and Supplementary Figure S3), although the number of HSR1 signals varied in *C. sativa* (Supplementary Figures S3C, S5). The overall comparative distribution of CS-1, HSR1, and HJSR satellite further suggests intergenomic changes during the species divergence (Figure 2).

**Figure 2.**
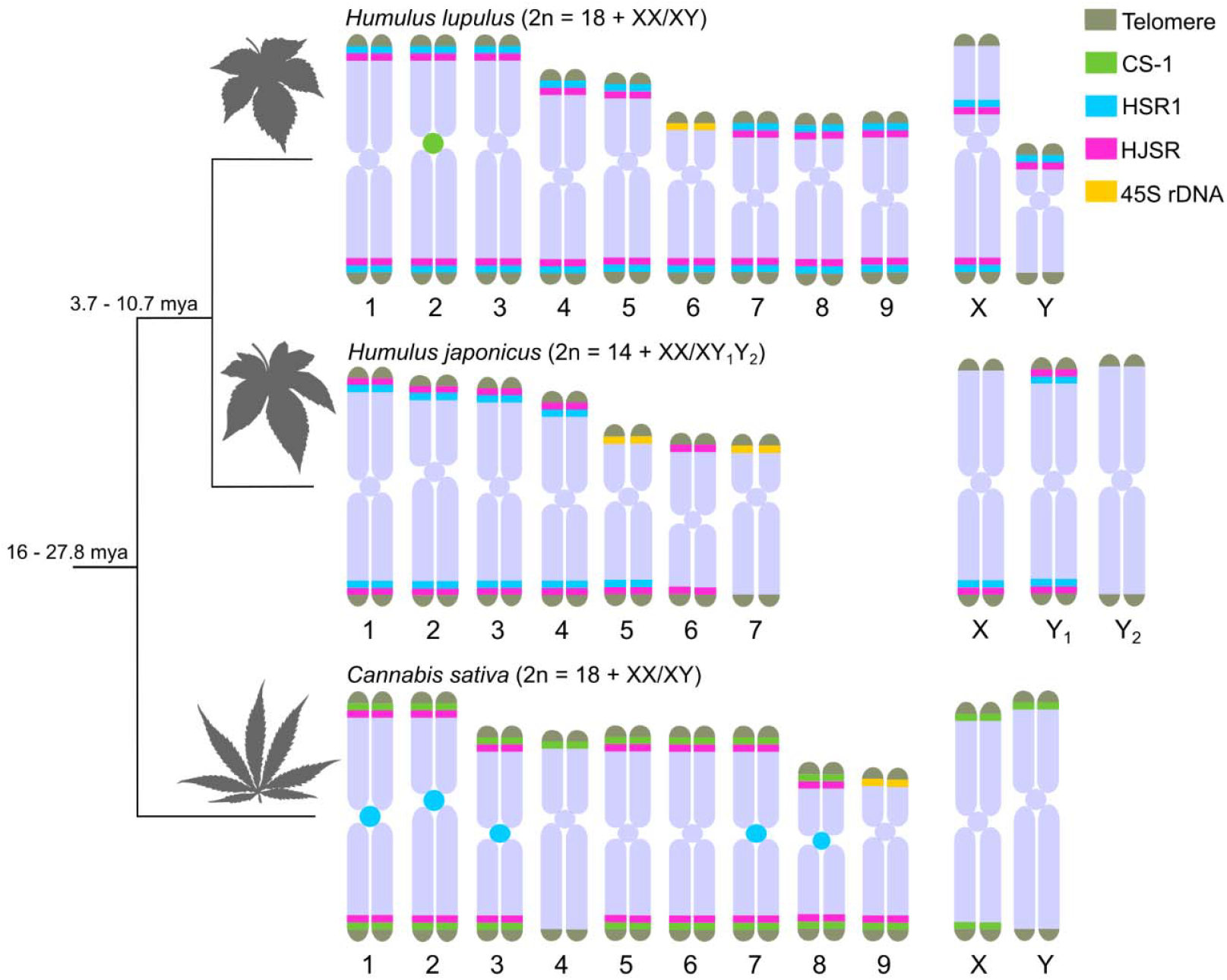
Idiograms illustrating chromosomal distribution of major satellites in *Humulus lupulus, H. japonicus*, and *Cannabis sativa*. The positions of telomere (grey), CS-1 (green), HSR1 (cyan), HJSR (magenta), and 45S rDNA (yellow) repeats were derived based on the FISH physical localization (Figure 1 and Supplementary Figures S4-S5). The relative relations among these species and estimated time of divergence were summarized according to Jin et al. (2020); McPartland (2018); Murakami (2000); Murakami et al. (2006); Padgitt-Cobb et al. (2023). Note that the presence of HSR1 and HJSR in *C. sativa* suggests a shared pool of satellite DNA sequences originating from their last common ancestor.

### 3.2. Characterization and origin of major satellites in *Cannabaceae*

We identify three satellites CS-1, HSR1, and HJSR within one supercluster family in the genomes of *Cannabis* and *Humulus* species using the RepeatExplorer2 pipeline (Novák et al., 2010, 2013). Based on this analysis, we extracted consensus sequence and determined their abundance in the genomes from sequencing reads (summarized in Table S). The monomer unit lengths for these satellites range from 370 bp for CS-1, 380 bp for HJSR, to 384 bp for HSR1. Sequence comparison revealed relatively high sequence similarity between HSR1 and CS-1 (74.3%) and HSR1 and HJSR (73.9%; Supplementary Figure SXY).

Using the same primers designed for each major satellite in the three species gave rise to species-specific ladder patterns for the CS-1 satellites in *Humulus* species (Supplementary Figure S2). The expected band size of 370 bp for the monomer unit was obtained exclusively in *C. sativa*. In contrast, the amplification of the HSR1 and HJSR satellites was consistent across species. The monomer length was 380 bp (Supplementary Figure S2). PCR products for each satellite and species were cloned and sequenced. We analyzed a total of 35 CS-1 sequences (23 from *C. sativa*, 5 from *H. lupulus*, and 7 from *H. japonicus*), 46 HSR1 sequences (15 from *C. sativa*, 14 from *H. lupulus*, and 17 from *H. japonicus*), and 36 HJSR sequences (11 from *C. sativa*, 14 from *H. lupulus*, and 11 from *H. japonicus*). A comparison of satellite sequences cloned from each species supported low interspecific variability and their classification into one supercluster family, in line with physical localization on metaphase chromosomes (Supplementary Figure S6). To determine the relationships among these satellites in studied species, we conducted a phylogenetic analysis of individual sequence clusters (Figure 3, Supplementary Figure S7-S8). The CS-1 amplified in *C. sativa* and *H. japonicus* grouped together, while those amplified in *H. lupulus* formed a separate clade (Figure 3). Although the sequences from *C. sativa* and *H. japonicus* are more closely related, no visible CS-1 signal was detected on the chromosomes of *H. japonicus* (Figure 1B), meaning low enrichment of these satellite units to be identified by FISH. In contrast, CS-1 was localized in the subtelomeric regions of *C. sativa* chromosomes. HSR1 and HJSR sequences tended to cluster together, and no unique species-specific clades between species were identified (Supplementary Figure S7-S8), representing low divergence between individual subunits, supported by satellite position in the subtelomeric region. In *C. sativa*, HSR1 was found in pericentromeric positions on five autosome pairs (Figure 1C). From all three satellite arrays, only the HJSR satellite displayed a conserved subtelomeric position on chromosomes across all three species (Figure 1D) again with no single clear and specific clade.

**Figure 3.**
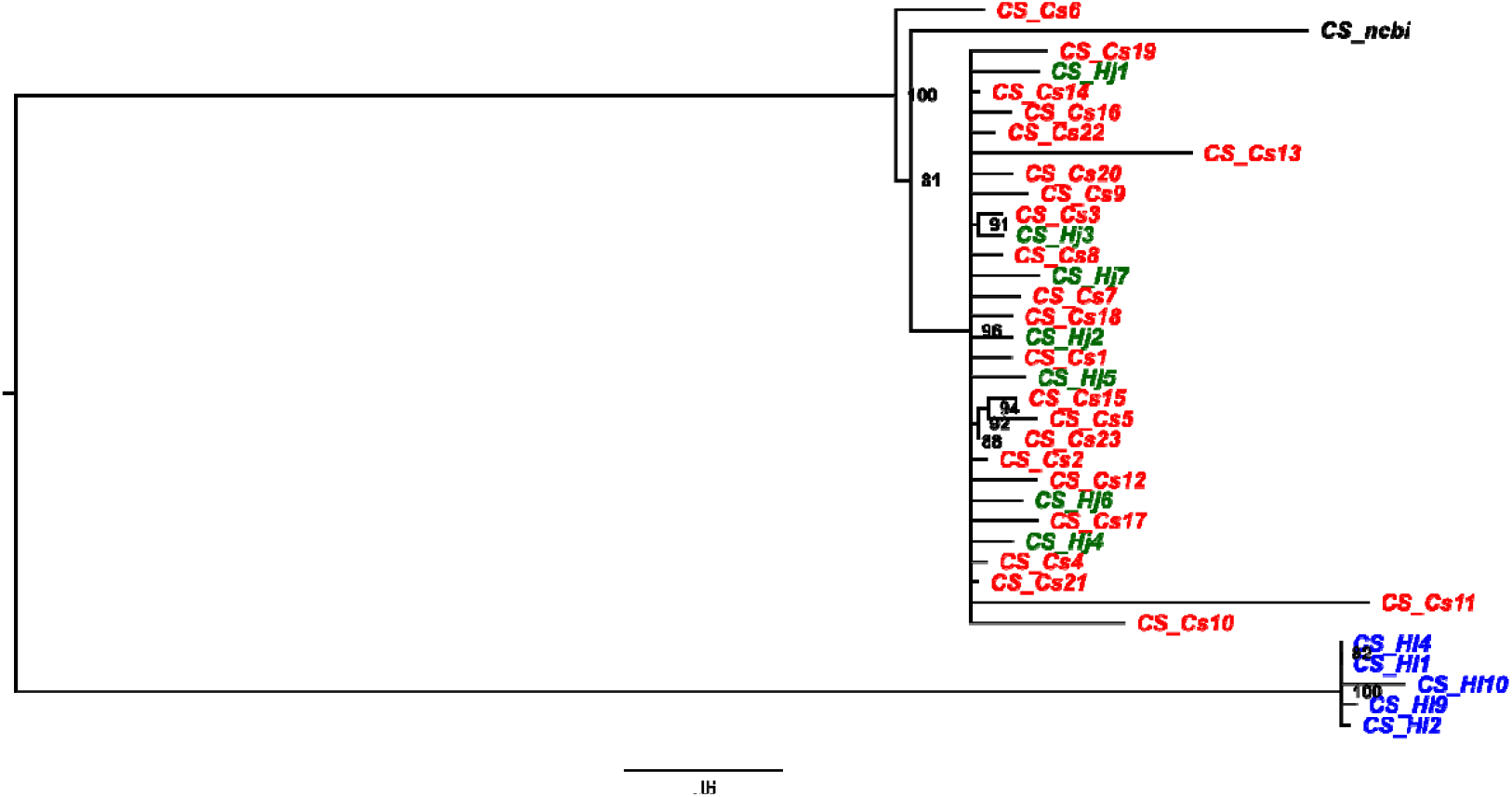
The phylogenetic relationships of sequences from *C. sativa, H. lupulus*, and *H. japonicus* amplified using primers designed for the CS-1 satellite. Note the separate clade for sequences from *H. lupulus*.

## 4. Discussion

Satellite DNA plays a crucial role in the organization of plant chromosomes and has critical implications for evolutionary, genetic, and taxonomic research. Analyzing repetitive DNA divergence enables the exploration of evolutionary relationships among plant species (Yoong Lim et al., 2006; Koo et al., 2011). Despite the progress in characterizing the satDNA content within *Humulus* species, the organization, location and the extent to which these sequences are shared or diverged among the related species of the *Cannabaceae* family remain unknown. In this study, we examined the molecular and cytogenetic characteristics and phylogenetic relationships of superfamily CS-1, HSR1, and HJSR in *C. sativa, H. lupulus*, and *H. japonicus*. According to the library hypothesis, related taxa share a library of different conserved satellite DNA units (different satellite DNA families, monomer variants, and subfamilies within a satellite DNA family). The satDNA sequences may be differentially amplified in each taxon with the subsequent replacement of one sequence variant by another in different species or populations (Plohl et al., 2012). Similarly, the evolution of the HSR1 and HJSR satellites is likely to be explained by such a process. The presence of both satellites in *C. sativa* suggests that they originated from a common ancestor, predating the speciation and split of *C. sativa* and *H. lupulus*. Although the number of repeat sequences varies significantly between species, differences in subcluster abundance likely result from species-specific amplification processes. (Supplementary Figures S7-S8; Plohl et al., 2008, 2012). In contrast, the CS-1 satellite appears more diverse than the HSR and HJSR satellites (Figure 3). Correspondingly, the satellite RAE180 in *R. acetosa* has undergone distinct pattern of accumulation on autosomes and sex chromosomes. It is supposed that RAE180 originated before the split between *Rumex* species with XX/XY and XX/XY_1_Y_2_ sex chromosome systems (Cuñado et al., 2007; Navajas-Pérez et al., 2009b, 2009a; Quesada Del Bosque et al., 2011). Since *Rumex* species contain so far, the highest number of satellites on the sex chromosomes in plants (Jesionek et al., 2021), satellite RAE180 was hypothesized to be present in an ancestral genome in various monomer variants from which novel tandem arrays could later be amplified (Navajas-Pérez et al., 2009a). Such divergence was assessed in *R. hastatulus* Texas and North Carolina cytotypes, *R. acetosella* and *R. acetosa* (Navajas-Pérez et al., 2009a; Quesada Del Bosque et al., 2011), supporting again library hypothesis as shown for supercluster family in this study.

In *H. lupulus* we obtained distinct sequences with pericentromeric localization on one chromosome pair (Figure 1B). Although we confirmed the presence of CS-1 in the *H. japonicus* genome, this satellite was not detected on the metaphase chromosome using standard FISH approach, even with low stringency and higher sensitivity. We suggest that the absence of CS-1 positive loci on *H. japonicus* chromosomes rather indicate its low abundance in the genome and low tandem organization, similar to the RAE180 satellite in *Rumex acetosella* (Cuñado et al., 2007). These findings are consistent with the large phylogenetic distance between *Cannabis* and *Humulus* genera and suggest distinct evolutionary trajectories for satellites in the lineages leading to *H. lupulus* and *H. japonicus*. SatDNA evolves rapidly through unequal crossing-over, replication slippage or mutation (Plohl et al., 2012). Amplification of satDNA may accompany karyotype rearrangements in plant species and chromosomal position of some satDNA family varies between related species (Navajas-Pérez et al., 2009a). In our study, we observed that the subtelomeric localization of the three major satellites was not conserved and showed the distinct distribution pattern across the studied species. Interspecific differences were evident in their chromosomal positions and number of foci. Despite these differences, subtelomeric localization was predominant. The *H. lupulus* HSR1 satellite was localized to the (peri)centromeric regions of *C. sativa*. Similarly, CS-1 was found exclusively in the (peri)centromeric region of two autosomes in *H. lupulus* (summarized in Figure 2 and Supplementary Table S3). This satellite localization (Figure 1C) supports the hypothesis of large-scale genomic reorganization during the evolution in *Humulus* (Akagi et al., 2024). This pattern is comparable to Cl12 in XY and XYY cytotypes of *R. hastatulus*, which during karyotype rearrangement was relocated from the (peri)centromeric regions and accumulated in subtelomeric regions of some autosomes and sex chromosomes (Sacchi et al., 2024). Despite chromosomal rearrangements, including autosome and X-autosome fusions that occurred during the evolution in *H. japonicus* (Akagi et al., 2024), leading to a reduced chromosome number and the formation of X and both Y chromosomes, telomeric signals were consistently observed only at the terminal regions of all chromosomes.

In summary, the analysis of the *Cannabaceae* supercluster satellite superfamily reveals a complex evolutionary pattern during the speciation of *H. lupulus* and *H. japonicus*. While differing in abundance, the sequence structure of CS-1, HSR1, and HJSR remains largely conserved and their chromosomal localization has contributed to species-specific sequence evolution. This divergence may have played a role in accelerating speciation, as seen in the distinct positioning of CS-1 and HSR1 arrays in *Cannabis sativa*. With the availability of long-read sequencing data, a comprehensive analysis of the full organizational structure of this supercluster family will become possible, facilitating the identification of new sequence motifs.

## Data availability statement

All datasets generated for this study are included in the article/Supplementary Material.

## Author contributions

VB, LH, and RH planned and designed research. LH, VB, and RČ performed experiments. JB performed phylogenetic analysis. LH and VB wrote the main text of the manuscript. JP provided plants of *H. lupulus*. All authors read and approved the final version of the manuscript.

## Acknowledgements

This work was supported by the Czech Science Foundation grant No. 22-00301S.

