## Supplemental Files for "Contrasting patterns of subtelomeric satellite superfamily in the *Cannabaceae* family"

| **Primer name** | **Orientation** | **Sequence (3´- 5´)** | **GenBank Accession** | **Reference** |
| --- | --- | --- | --- | --- |
| OPJ9 | Forward | ACAGAGTACAACTCAGAAACAAACC | KX688593.1 | Polley et al. 1997 |
|  | Reverse | AAGGTCGCACAATGACCG |  |  |
| MADC2 | Forward | GTGACGTAGGTAGAGTTGAA |  | Mandolino et al. 1999 |
|  | Reverse | GTGACGTAGGCTATGAGAG |  |  |
| HSR1 | Forward | CCCTCTGGTGAATTGGAGAT | GU831574.1 | Divashuk et al. 2011 |
|  | Reverse | CCTTTCAGAAATCTTCGATTTCTCTA |  |  |
| HJSR | Forward | GCGATGAGCTTACTATTTCTTCCA | GU831573.1 | Alexandrov et al. 2012 |
|  | Reverse | CGATGGGATAACCGAAGAATTGG |  |  |
| CS-1 | Forward | GGTACCACTATGAGAAATGTGAG | JX402748.2 | Divashuk et al. 2014 |
|  | Reverse | CCTTTGTGAAATGTGGCCCGG |  |  |
| CS-1 (for *H*. *japonicus*) | Forward | GGTACCACTATGAGAAATGTGAGA |  |  |
|  | Reverse | AAATGTGGCCCGGACTCATC |  |  |
| 45S rDNA | Forward | TGCCCGTTGCTCTGATGATT | AF223066.1 |  |
|  | Reverse | TCCACCAACTAAGAACGGCC |  |  |
| Telomere | Forward | TTTAGGGTTTAGGGTTTAGGGTTTAGGGTTTAGGG |  | Ljdo et al. 1991 |
|  | Reverse | CCCTAAACCCTAAACCCTAAACCCTAAACCCTAAA |  |  |

Table S1 Sequences of primers used for PCR amplification and preparation of DNA probes for FISH analysis.

| **Hybridization mixture**  **Stringency** | 77% | 68% |
| --- | --- | --- |
|  | **Volume**  (µl) | **Volume**  (µl) |
| **100% formamide** | 10 | 7 |
| **50% dextran sulphate** | 4 | 4 |
| **20x SSC** | 2 | 2 |
| **HSR1/HJSR/CS-1** | 1 | 1 |
| **45S rDNA** | 1 | 1 |
| **Water** | 2 | 5 |
| **Total volume** | 20 | 20 |

Table S2 The composition of hybridization mixture for selected stringency conditions used in the study.

| **Tandem repeats** | **PCR** | | | **Chromosomal localization** | | |
| --- | --- | --- | --- | --- | --- | --- |
|  | *C*. *sativa* | *H*. *lupulus* | *H*. *japonicus* | *C*. *sativa* | *H*. *lupulus* | *H*. *japonicus* |
| CS-1 | + | + (different ladder pattern) | + (different ladder pattern) | +/subtelomere | +/(peri)centromere | +/subtelomere |
| HSR1 | + | + | + | +/(peri)centromere | +/subtelomere | +/subtelomere |
| HJSR | + | + | + | - | +/subtelomere | +/subtelomere |

**Table S3** A comprehensive overview and chromosomal localization of the three satellite sequences CS-1, HSR, and HJSR in studied species, *C*. *sativa*, *H*. *lupulus*, and *H*. *japonicus*.

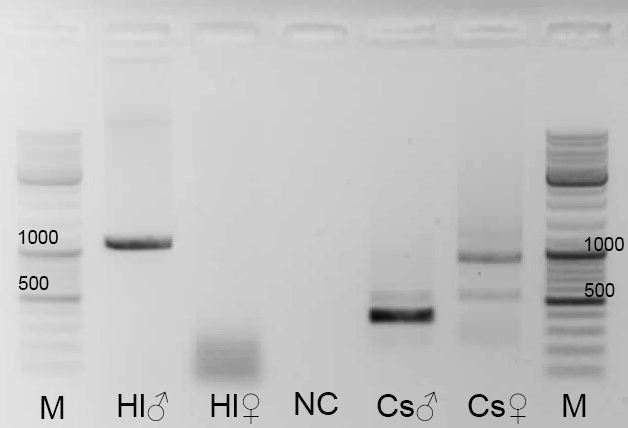

**Figure S1** The PCR-based sex determination in *Humulus lupulus* and *Cannabis sativa*. In *H*. *lupulus* male (Hl ♂), the OPJ9 primers amplified a male-specific region of approximately 1.15 kb, which was absent in the female *H*. *lupulus* (Hl ♀). Similarly, in *C*. *sativa* male (Cs **♂**), the male-associated MADC2 marker, with an estimated length of 390 bp, was clearly amplified only in male individual(s) and was not detected in *C*. *sativa* female (Cs♀). PCR products were separated in 1.5% agarose gel stained with ethidium bromide (EtBr). M – DNA ladder, NC – negative control.

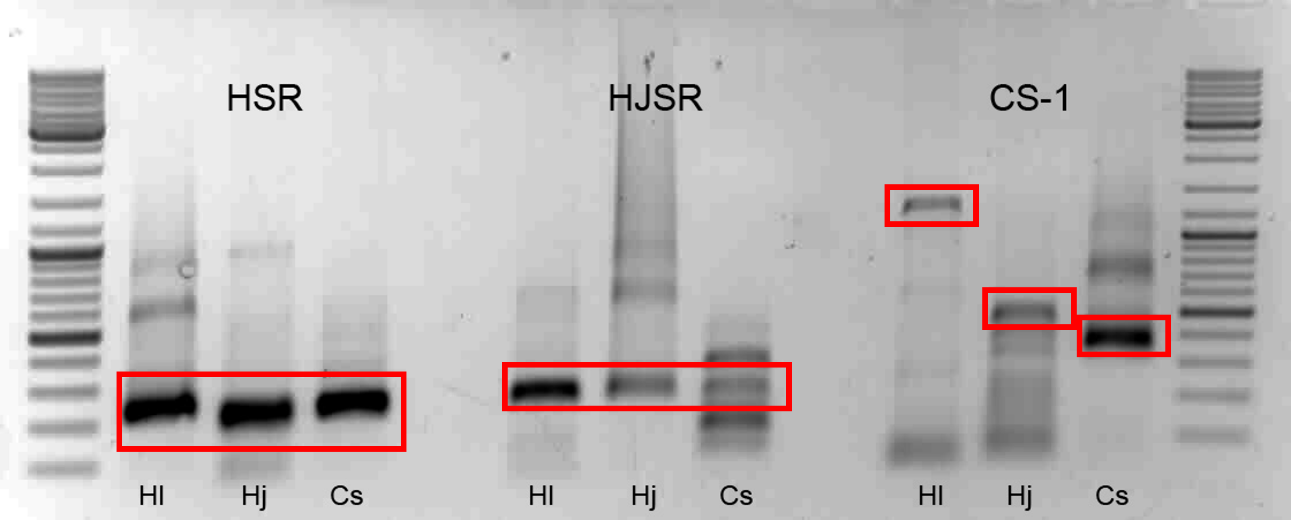

**Figure S2** PCR-based amplification of HSR1, HJSR and CS-1 subtelomeric satellites. The patterns of HSR1 and HJSR are consistent in *H*. *lupulus* (Hl), *H*. *japonicus* (Hj), and *C*. *sativa* (Cs), using the same pair of primers. The CS-1 sequence exhibits a distinct pattern for each species. PCR products were separated in 1.5% agarose gel with EtBr staining. The red rectangles indicate selected DNA fragments, for each satellite and for each species.

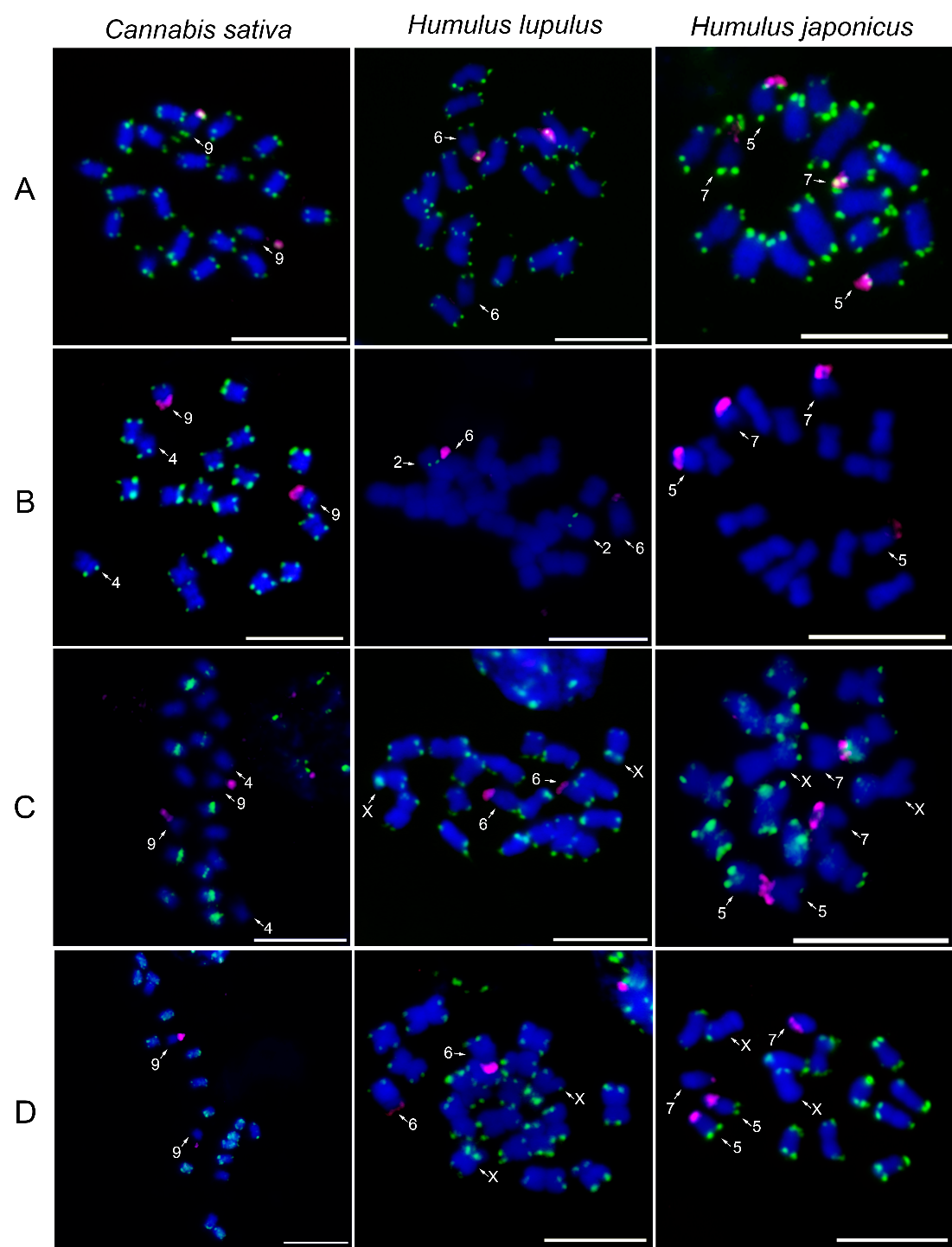

**Figure S3** Chromosomal distribution of major satellite repeats in studied species. The localization of (A) telomeric sequence motifs (TTTAGGG, green), (B) CS-1 satellite (green), (C) HSR1 satellite (green), (D) HJSR satellite (green), and 45S rDNA (magenta) on female mitotic metaphase chromosomes of *Cannabis* *sativa*, *Humulus* *lupus*, and *H*. *japonicus*. Note the reduced number of CS-1 positive regions in *H*. *lupulus* and missing localization of the same repeats in *H*. *japonicus* (B) for both used stringencies (68 % and 77%). Interestingly, HSR1 is localized within the (peri)centromeres in *C*. *sativa*, while displaying subtelomeric position in *H*. *lupulus* and *H*. *japonicus* (C). Mitotic chromosomes were counterstained with DAPI. The arrows indicate differentiated autosomes and sex chromosomes. Scale bar = 10 µm.

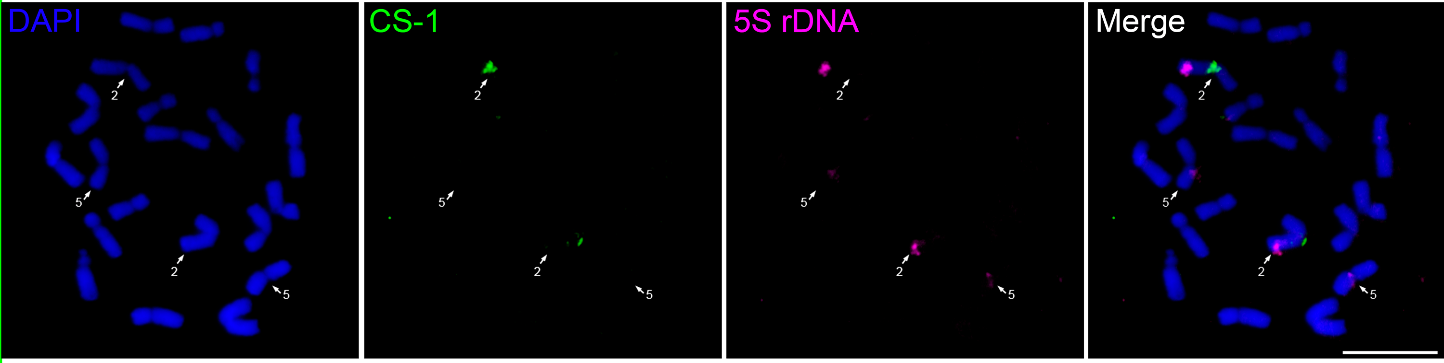

**Figure S4** Localization of CS-1 satellite repeat and 5S rDNA in male *Humulus lupulus*. The distribution of 5S rDNA (magenta) is consistent with the karyotype analysis described in Karlov et al. (2003). The 5S rDNA is localized in the subtelomeric region of chromosome 2 and the pericentromeric region of chromosome 5. Notably, both 5S rDNA and CS-1 satellite (green) are localized on chromosome 2. Mitotic chromosomes were counterstained with DAPI. Scale bar = 10 µm.

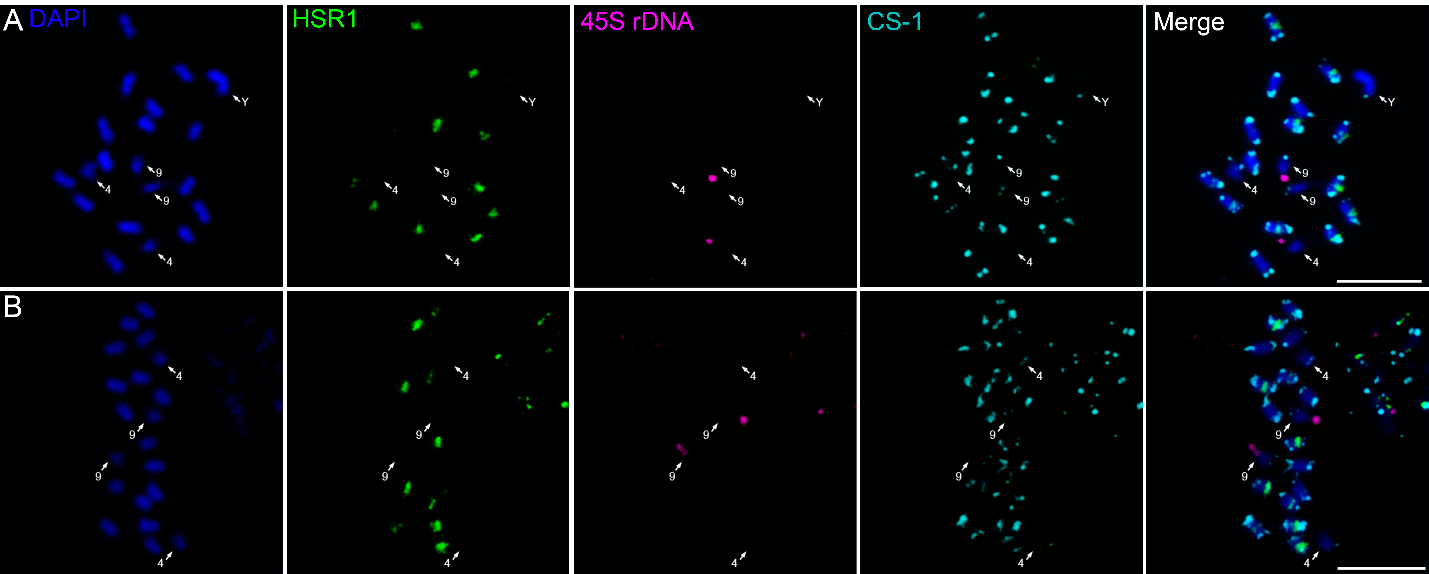

**Figure S5** The colocalization of selected satellite repeats in *C*. *sativa*. The distribution of HSR1 (green) and CS-1 (cyan) satellite repeats on metaphase chromosomes is shown for (A) male and (B) female *Cannabis* *sativa*. Ribosomal 45S rDNA (magenta) was used as a reference marker to identify chromosome 9. HSR1 exhibits (peri)centromeric signals on, the ten chromosomes in (A) male and on eleven chromosomes in (B) female. Mitotic chromosomes were counterstained with DAPI. Scale bar = 10 µm.

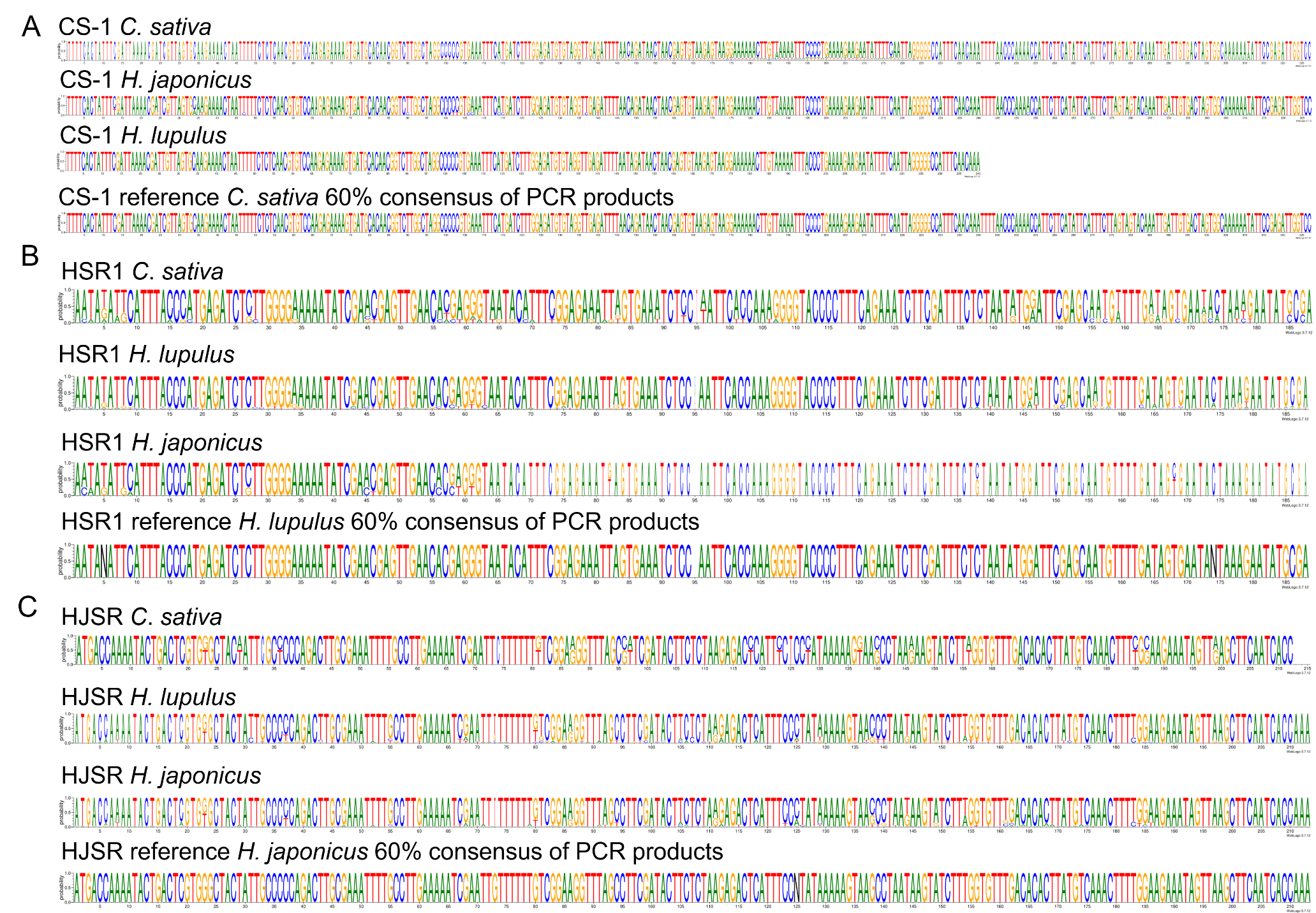

**Figure S6** Sequence logos and variability of major satellite repeats in studied species. The sequence variability of the major satellite repeats (A) CS-1, (B) HSR1, and (C) HJSR is shown for *Cannabis* *sativa*, *Humulus* *lupus*, and *H*. *japonicus*. Reference sequences were constructed from the consensus of cloned and isolated fragments, representing a 60% consensus sequence for each tandem repeat. The height of each letter in the sequence logo reflects its observed frequency at a particular position in the alignment. Notably, the length of the CS-1 repeat sequence varies among studied species, being overall shorter in *H*. *lupulus* than in *H*. *japonicus* and *C*. *sativa*.

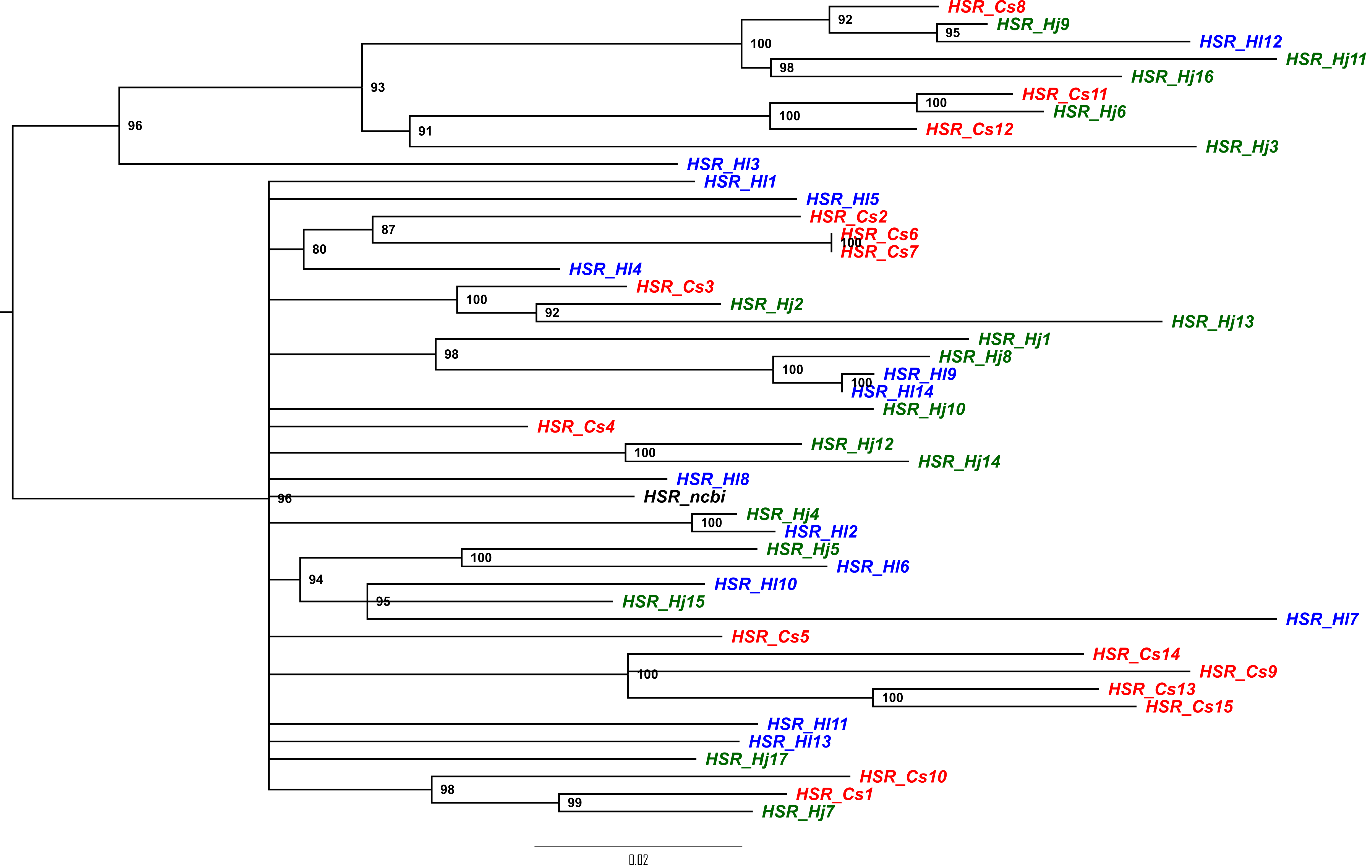

**Figure S7** Phylogenetic tree of HSR sequences from *C*. *sativa*, *H*. *lupulus*, and *H*. *japonicus*.

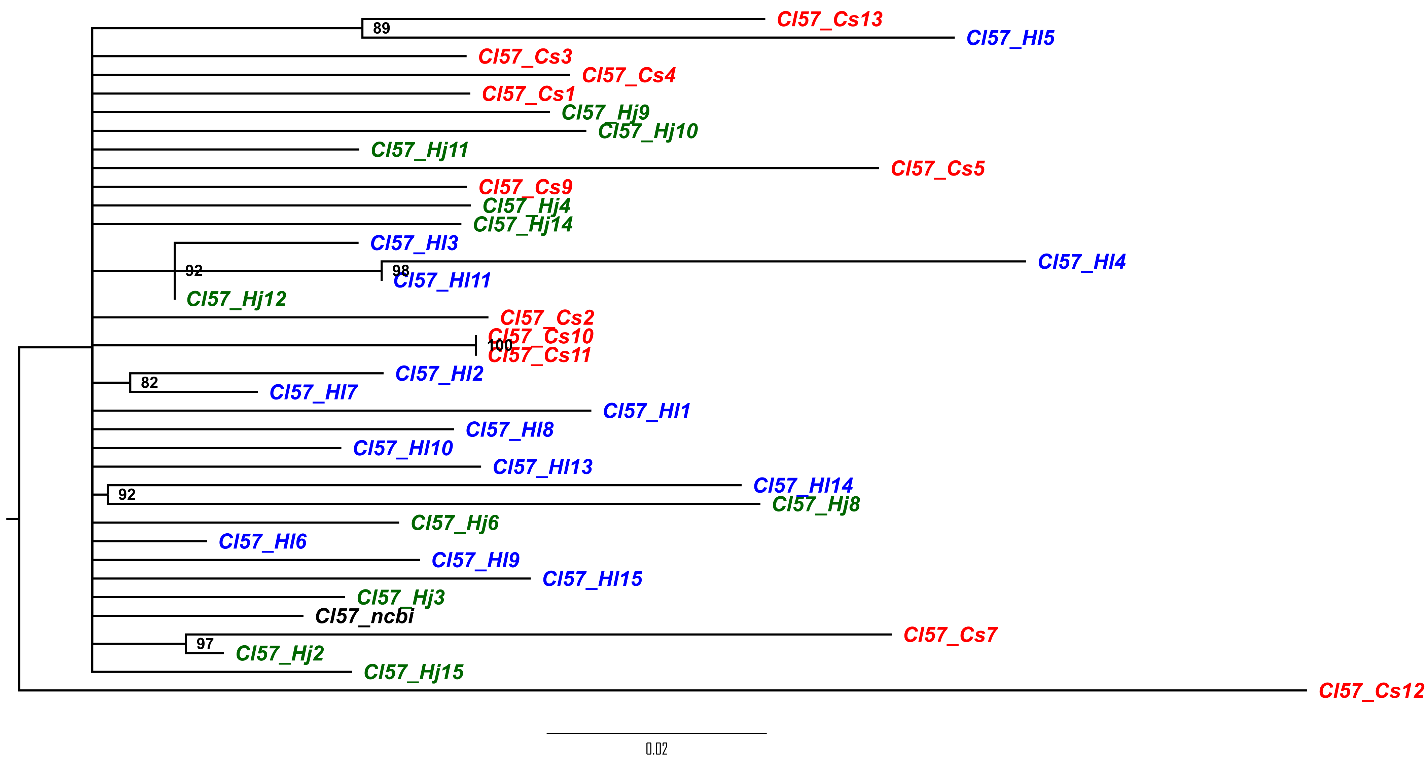

**Figure S8** Phylogenetic tree of HJSR sequences from *C*. *sativa*, *H*. *lupulus*, and *H*. *japonicus*.
